# Further evidence of increased human *Herpesvirus* in Alzheimer’s disease

**DOI:** 10.1101/858050

**Authors:** Ben Readhead, Jean-Vianney Haure-Mirande, Michelle E. Ehrlich, Sam Gandy, Joel T. Dudley

## Abstract

The recent coordination of scientific and political efforts within the field of Alzheimer’s disease (AD) research coupled with the rapid evolution of molecular profiling technologies has enabled the generation of unprecedented repositories of biological sequence data available for investigation by the scientific community. Next-generation sequences (NGS) represent complex readouts of cellular transcriptomic and genomic states, and hold the potential to illuminate disease biology via unexpected applications of collected data. We recently reported a multiomic study of the Alzheimer’s associated brain virome, leveraging NGS data to understand the potential relevance of viral activity to AD. We observed increased abundance of human herpesvirus 6A (HHV-6A), HHV-7, and herpes simplex virus 1 (HSV-1) genomes in banked postmortem brains from subjects with AD compared with controls. These results were replicated in two additional, independent and geographically dispersed cohorts. Although these findings might be of general interest to the field, such analyses present novel methodological challenges that have not been well characterized. The viral abundances that we detected were generally quite low, and, in many samples, no virome was detected. We sought to understand whether such sparsity might be associated with the spurious detection of differential abundance. Herein, we employ a simulation approach, which indicates that sparsity in abundance actually biases against the detection of significant differences between AD and control subjects. We also report new results based on previously unpublished whole genome sequences that show increases in both abundance and prevalence of HHV-6A in AD, consistent with our previously reported results. These findings, as well as reports from many other groups employing a variety of approaches, support the dedication of a coordinated effort toward a comprehensive characterization of the microbiome of the brain affected by AD. Such studies should advance our understanding of the potential clinical relevance of these observations.

## Introduction

The potential influence of pathogenic microbes on the risk of developing or accelerating the progress of Alzheimer’s disease (AD) was first proposed by Alois Alzheimer^1^ and Oskar Fischer^2^ well over a century ago. Since this time, and particularly over the past four decades, hundreds of scientific papers have emerged linking the activity of diverse microbial species with different facets of AD pathophysiology.

These reports have accelerated even further recently, including reports of *Herpesviridae* induced seeding of Aβ fibrillation in transgenic AD mouse and 3D organoid model systems^3^. In a population-based cohort study in Taiwan, Tzeng *et al^4^* observed an increased dementia risk following HSV-1 infection. These authors reported that the dementia risk in this cohort post HSV-1 infection appeared to be almost completely abolished by antiviral use. Lövheim *et al^5^* described an apparent association between the HSV-1 carrier state and episodic memory decline, particularly within *APOE ε4* carriers. Lindman *et al*^6^ reported that *APOE ε4* carriage interacts with additional AD genetic risk factors to potentiate the impact of HSV-1 on the development of AD. Recently, Rizzo *et al^7^* reported that Natural Killer (NK) cell receptor haplotype KIR2DS2/KIR2DL2/HLA-C1 is apparently associated with increased risk for developing AD and that NK cells expressing this combination of receptors demonstrated a reduced ability to clear HHV-6A infected T-cells because NK cell cytotoxicity was impaired. The authors also describe increased *APOE* expression in HHV-6A infected T-cells. Our own study in this area described an increased abundance of HHV-6A, HHV-7, and HSV-1 in the brains of subjects with AD, an observation that persisted in additional patient cohorts, and that was strengthened during a meta-analysis across all available samples^8^.

Throughout our study, we adopted a multi-stage viral mapping approach which includes a preliminary mapping of RNA sequences to a viral database in order to identify candidate viral reads, followed by filtering of human sequences and bacterial sequences, trimming of low quality reads, and a final mapping to a database of viral 31mers where each 31mer was selected for its specificity to a single virus, and for its absence from the reference human genome. Our motivation for using this approach was based on initial observations within the Mount Sinai Brain Bank (MSBB) of an increased abundance of HHV-6A, HHV-6B, and HHV-7 using a more straightforward subtractive mapping approach. These three *Roseoloviruses* share substantial sequence homology, as well as including flanking telomeric repeat regions that are not readily distinguishable from human telomeric repeats using short read sequences. The goal of our approach was to enable us to distinguish between homologous viruses and to minimize misclassification of human-derived reads as viral, we thus optimized for mapping specificity, rather than sensitivity.

This approach introduces certain limitations, including what are likely to be heterogeneous effects on the power to detect group level differences in viral abundance. This would almost certainly reduce our sensitivity for detecting viruses with high homology either to other viruses or to human sequences. In addition, due to the combination of filtering, and mapping to the viral 31mer database, the approach will systematically underestimate true viral abundance. In situations where viral abundance may only be moderate (yet still biologically interesting), this underestimation coupled with the possibility that significant fractions of the population under consideration might not have any detectable virus introduces novel challenges to differential expression methodologies that have been developed and optimized for characterizing endogenously expressed transcripts.

Our strategy for mitigating such biases was to apply diverse, integrative analytical approaches for understanding the relevance of viruses to AD rather than relying on any single finding. While observations of differential abundance between AD and control subjects were informative and served as a conceptual starting point for our investigations, we found substantial value in subsequent integrative analyses, including the association of viral abundance with clinical and neuropathological traits, integration of DNA sequence data to identify viral quantitative trait loci (vQTL), and construction of virus-host regulatory networks. For example, throughout our study we observed a surprisingly high prevalence of certain viral species (e.g. Hepatitis C virus, Variola Virus) which almost certainly represent false positives based on prior knowledge from clinical and epidemiological studies. However, we did not see observe strong relationships between these species and AD-relevant clinico-pathological traits, vQTL, virus-host gene regulatory networks, and thus these species were not emphasized in our report. Thus, while our approach for metagenomics classification is not well suited for estimating population prevalence for certain viral species, the integration of additional multiomic data layers does allow us to identify species that associate with AD from multiple, diverse perspectives.

Here, we review the quantification of viruses that we reported as differentially abundant in the AD brain (HHV-6A, and HHV-7)^8^. Using a simulation approach, we show here that while these viruses were generally detected at low levels, the adopted methodology is capable of detecting potentially meaningful group level differences. We find that the structure of these data (including many samples without detected virus) actually biases *against* the detection of differential abundance, rather than potentially leading to spurious association with AD.

We also present new findings from whole genome sequences (WGS) from post-mortem brain tissue generated from 326 subjects in the MSBB, using a complementary viral quantification approach based on *de novo* viral contig assembly. These data replicate our original finding of increased HHV-6A abundance in AD, and show that in addition to being present at higher abundance in AD brain, HHV-6A is also present at higher prevalence in AD brain when compared with brains from matched controls.

This study builds upon our previous report^8^, demonstrating the potential opportunities for detection of non-human sequences that are inadvertently captured in human NGS data. We also note the heterogeneity in viral signal captured via different sequencing modalities. The re-emergence of HHV-6A in this new dataset provides an encouraging consistency in viral signal. In combination with the findings of many other investigators^3, 4, 6, 7, 13, 15-21^ these new data also support the utility of a systematic, coordinated effort to characterize the brain microbiome in AD.

## Results

### Low viral abundance and sparsity of HHV-6A and HHV-7 in RNA-sequence data biases against the detection of differences between AD and normal subjects

In our report^8^, we adopted a differential expression analysis paradigm to investigate differences in viral abundance between AD and controls. A common challenge in such comparative transcriptomics is the issue of filtering features (usually endogenous genes, though in our case, viruses, or viral genes) on the basis of their detection in a sufficient subset of samples. In the context of viral sequences, this can mean that large fractions of the samples might comprise “zero-counts” (where no virus was detected). As this fraction increases, the statistical assumptions underlying the methodologies may not be fulfilled, and potentially compromise the ability to accurately define group-level differences.

We evaluated RNA-sequences generated from subjects within the MSBB across four brain regions, and found that the proportion of zero counts varies substantially according to brain region and viral species (**Table 1)**. In the regions where we reported differential abundance of HHV-6A, and HHV-7 (anterior prefrontal cortex: APFC, and superior temporal gyrus: STG, **Figure 1**), approximately 30-35% of samples had no reads mapping to these viruses (zero-counts).

**Table 1.** Detection of HHV-6A and HHV-7 in MSBB.

| Tissue | Virus | Diagnosis | Mean log2(CPM) | N(Zero Counts) | N(Samples) | Proportion Zero Counts |
| --- | --- | --- | --- | --- | --- | --- |
| APFC | HHV6A | AD Possible | -4.83 | 45 | 145 | 0.31 |
|  |  | Control | -5.07 | 26 | 68 | 0.38 |
|  | HHV7 | AD Possible | -4.77 | 43 | 145 | 0.30 |
|  |  | Control | -4.98 | 25 | 68 | 0.37 |
| STG | HHV6A | AD Possible | -4.29 | 16 | 67 | 0.24 |
|  |  | Control | -4.73 | 8 | 29 | 0.28 |
|  | HHV7 | AD Possible | -4.36 | 18 | 67 | 0.27 |
|  |  | Control | -4.86 | 13 | 29 | 0.45 |
| PHG | HHV6A | AD Possible | -5.48 | 46 | 71 | 0.65 |
|  |  | Control | -5.24 | 18 | 36 | 0.50 |
|  | HHV7 | AD Possible | -5.16 | 31 | 71 | 0.44 |
|  |  | Control | -4.89 | 12 | 36 | 0.33 |
| IFG | HHV6A | AD Possible | -5.25 | 68 | 138 | 0.49 |
|  |  | Control | -5.08 | 20 | 48 | 0.42 |
|  | HHV7 | AD Possible | -5.18 | 65 | 138 | 0.47 |
|  |  | Control | -5.07 | 21 | 48 | 0.44 |

**Figure 1.**
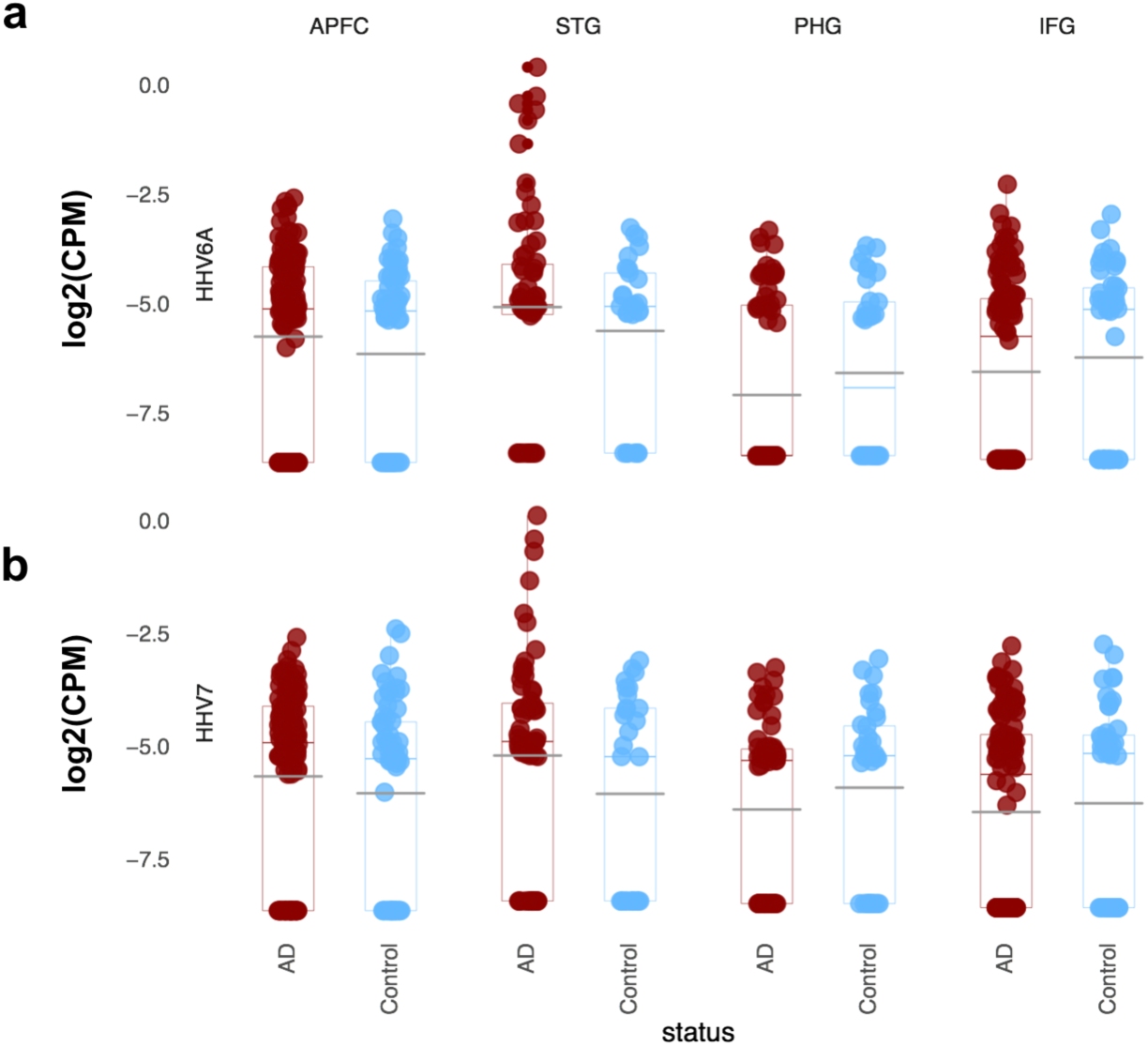
Abundance of Viral RNA in the MSBB. Multiregional comparison of **(a)** HHV-6A, and **(b)** HHV-7 abundance between AD versus controls, showing log2(CPM). A prior count of 0.1 was used to avoid taking the log of zero counts. Boxplots show interquartile range, with the horizontal colored bar denoting median, and gray denoting mean abundance.

We also noted differences between these fractions, and the fractions of zero counts in the parahippocampal gyrus (PHG), and inferior frontal gyrus (IFG) (regions where we did not detect differential abundance). In those regions, for approximately 45-50% of samples, we did not detect any reads mapping to HHV-6A, or HHV-7. Whether these differences are driven by biological or technical differences in samples from these regions is not clear, but this illustrates a critical element in our choice of methodology, through the use of Voom^9^. The Voom (variance modeling at the observational level) method models the mean-variance trend for count data, and generates a precision weight for each observation. Low read counts (which are associated with a higher variance) are assigned a correspondingly lower precision estimate, which is incorporated into subsequent linear modelling and differential abundance steps.

We sought to clarify what the impact of high zero-counts might be on the methodology used to detect differential viral abundance in our study, and whether this might lead to an inflation of p-values and drive spurious differences in HHV-6A, and HHV-7 between AD and control subjects. Overall, we observed that our application of this method in the context of detecting differences between groups, based on low count data, biases against the detection of significant differences. **Figure 2** summarizes the effect of progressively altering the read count of non-zero samples (for HHV-6A and HHV-7) to zero, and demonstrates the effect of this on differential abundance results obtained within samples from the MSBB APFC and STG (the regions where we initially observed differential abundance). For each virus, we extract the set of non-zero samples and then iterate progressively down this list, varying the read counts for the virus under consideration to zero, and performing differential abundance analysis as previously reported. To remove path-dependent effects that could arise from the order in which samples are zeroed, we performed 1000 permutations and report the median –log10(FDR) for each increment in sample zeroing, for each virus. As progressively larger fractions of samples are zeroed, there is a rapid drop in the differential abundance FDR for both HHV-6A, and HHV-7. These findings suggest that the evaluation of viral abundances in the context of high zero-counts actually biases against the detection of significant between-group differences. Despite our observations of an AD-associated increase of HHV-6A and HHV-7 being based on abundance counts that are lower than normally adopted in comparative transcriptomics of endogenous genes, these constraints appear to limit our power to detect group level differences, and correspondingly that this is not associated with an inflation of false positives.

**Figure 2.**
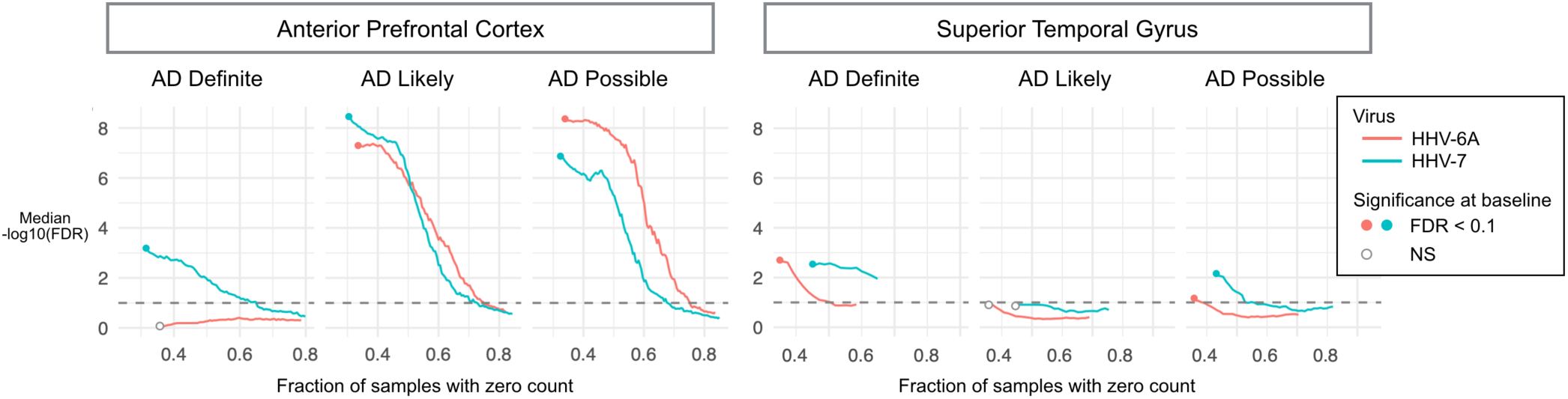
Reduced power to detect differences with increasing zero count. These graphs demonstrate the effects of progressively altering viral counts to zero on differential abundance analysis for HHV-6A and HHV-7 in the two brain regions of MSBB where differential abundance was originally reported. As progressively larger fractions of samples are artificially altered to zero, the detection of significant differences between groups declines rapidly, indicating that this approach is biased against the detection of differences, rather than the inflation of false positives. Median –log10(FDR) of differential abundance from 1000 permutations of each zeroing increment shown.

### Whole genome sequences from post-mortem brain tissue in MSBB also demonstrate increased abundance of HHV-6A in Alzheimer’s disease cases using an alternative quantification approach

Throughout our published study^8^, we utilized a multi-stage mapping approach, where candidate viral reads were mapped to a database of 31mers that had been selected on the basis of specificity to a single virus and without homology to the human reference genome. The goal of using this approach was to enable us to distinguish between homologous viruses and to minimize misclassification of human sequences as viral. This approach has the associated effect of underestimating viral abundance and likely contributes to the generally low viral read counts we reported.

In order to evaluate the veracity of our findings using a complementary approach, we applied an alternative viral quantification method to WGS that have been generated from STG tissue from 326 subjects within the MSBB (AD Definite: n=158, AD Likely: n=40, AD Possible: 39, Normal: n=89). These samples were not included in our original report. We adopted a contig clustering based approach (see Methods for detailed description) that comprises an initial alignment of WGS fastq files to a viral reference database, followed by removal of human reads, trimming of low quality reads, removal of likely bacterial reads, assembly of remaining reads into contigs, and formation of contig clusters that are associated to specific viral species using BLASTn. We retained contig clusters with length > 300 nucleotides, and a mean coverage > 5x for subsequent analysis. Virus level abundance for each sample was estimated as the sum of mapped reads for each contig cluster associated with a particular virus.

Across the 326 samples, we identified 768 contig clusters, mapping to 42 unique viruses (**Supplementary Data 1**), derived from 270 of the 326 (86%) subjects. The virus with the longest mean contig clusters detected was HHV-6A (7500 nucleotides), and HHV-6B (7144 nucleotides), which were each detected in 14 (4.3%), and 15 (4.6%) samples respectively.

We performed differential viral abundance analysis in a manner comparable with the approach previously described^8^ (see Methods). We observed a significantly increased abundance of HHV-6A in each of the three AD vs. Normal comparisons (**Figure 6a**), consistent with our previous report^8^. This difference was most robust in a comparison of AD Definite vs. Normal (FDR < 4e-5). We also detected an increased abundance of HHV-6B in one comparison (AD Definite vs. Normal) and an increased abundance of Torque Teno Midi Virus 2 (TTMV2) in two comparisons.

We found that a substantial component of the differences in abundance for these three viruses were driven by large differences in the fraction of AD cases and Normals with detectable virus (**Figure 3b, Table 2**), particularly for HHV-6A (AD Definite vs. Normal, Fisher’s exact test, P-value: 4.85e-3). We detected HHV-6A in 14 subjects overall, primarily in subjects classified as AD Definite, and did not detect it in any Normal subjects (AD Definite: n=12, AD Likely: n=1, AD Possible: n=1, Normal: n=0) (**Supplementary Table 2**). Although the fold-enrichments for TTMV2 (Obs/Exp: 3.22), and HHV-6B (Obs/Exp: 3.56) were positive, they did not reach nominal significance.

**Table 2.**
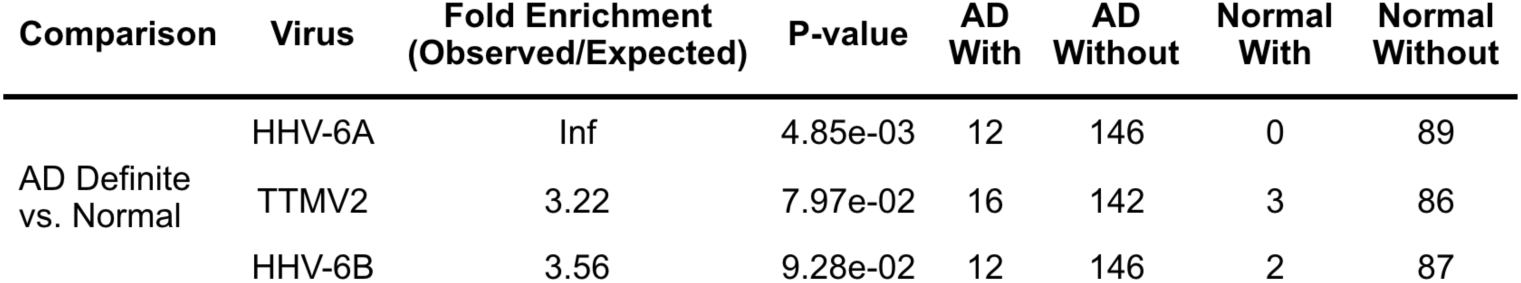
Differential prevalence of HHV-6A in AD.

**Figure 3.**
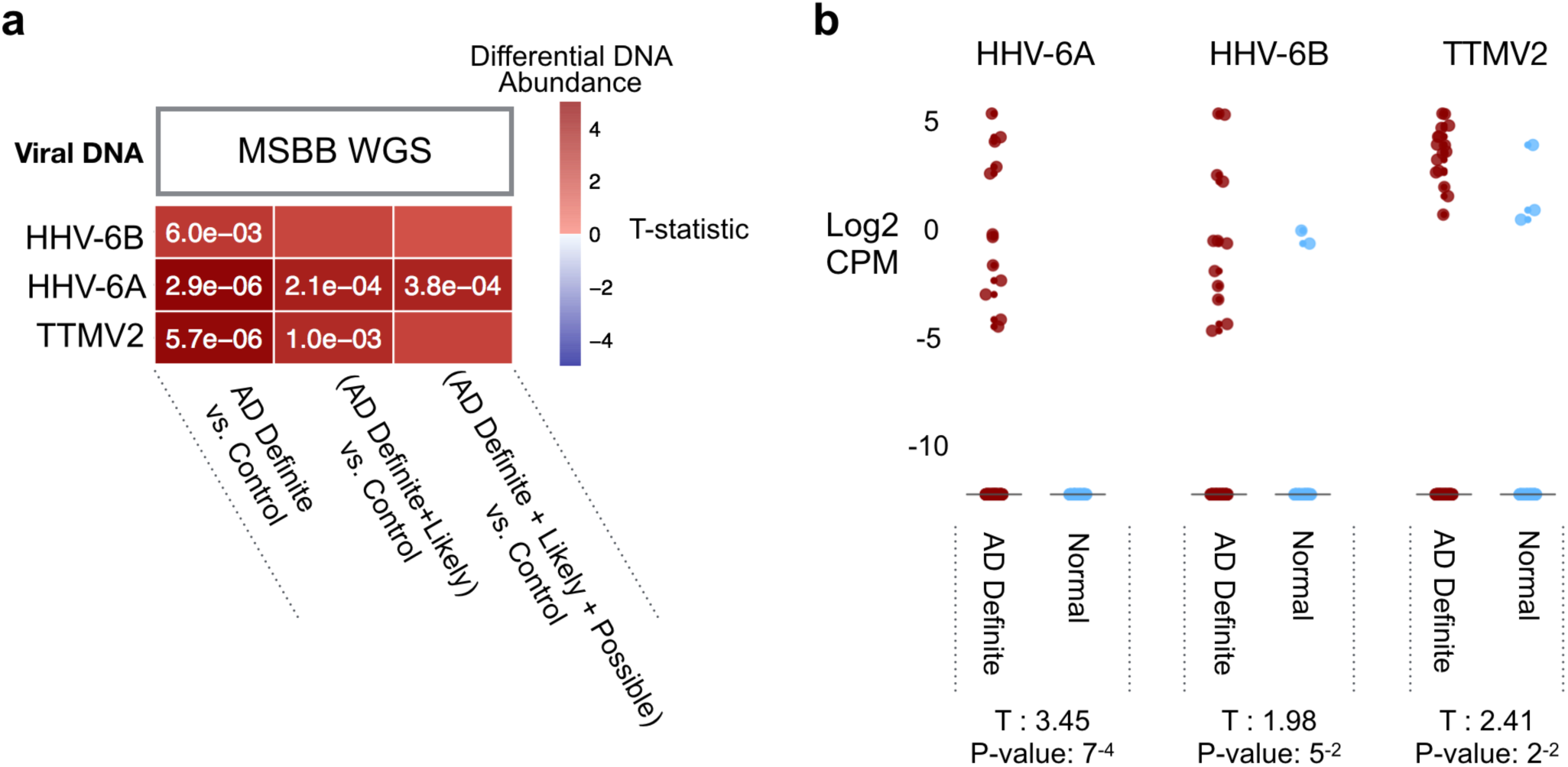
Differential abundance of viral DNA in whole genome sequences. Viral abundance estimated using a *de novo* contig assembly based approach applied to whole genome sequences generated from 326 samples within the MSBB cohort. **(a)** Heatmap summarizing results from Voom based analysis for viruses significantly (FDR < 0.1) differentially abundant in at least one comparison. P-values shown in cells with FDR < 0.1. **(b)** Boxplots of log2(CPM) for three viruses implicated in Voom based analysis. CPM: Counts per million.

Overall, the replication of a key finding (an increased abundance of HHV-6A in AD) using a conceptually distinct approach to viral quantification, applied to a novel molecular dataset that was generated from the same MSBB cohort we reported on originally, supports the robustness of our original observation.

### Viral presence associates with clinico-pathological traits of Alzheimer’s disease

To explore the potential relationships between the presence of the three viruses detected in our differential abundance analysis of the MSBB WGS data, we sought to determine whether there existed systematic differences in clinical dementia rating (CDR), mean amyloid plaque density, and Braak and Braak score (Braak score) when comparing subjects with, and without each virus (**Figure 4**). We observed that HHV-6A was most significantly positively associated with mean amyloid plaque density, and Braak score (an indicator of neurofibrillary tangle burden), and was nominally significantly associated with clinical dementia rating. Interestingly, TTMV2 was also significantly associated with each of these three traits, most robustly with CDR. We did not detect any significant associations between HHV-6B and these AD traits.

**Figure 4.**
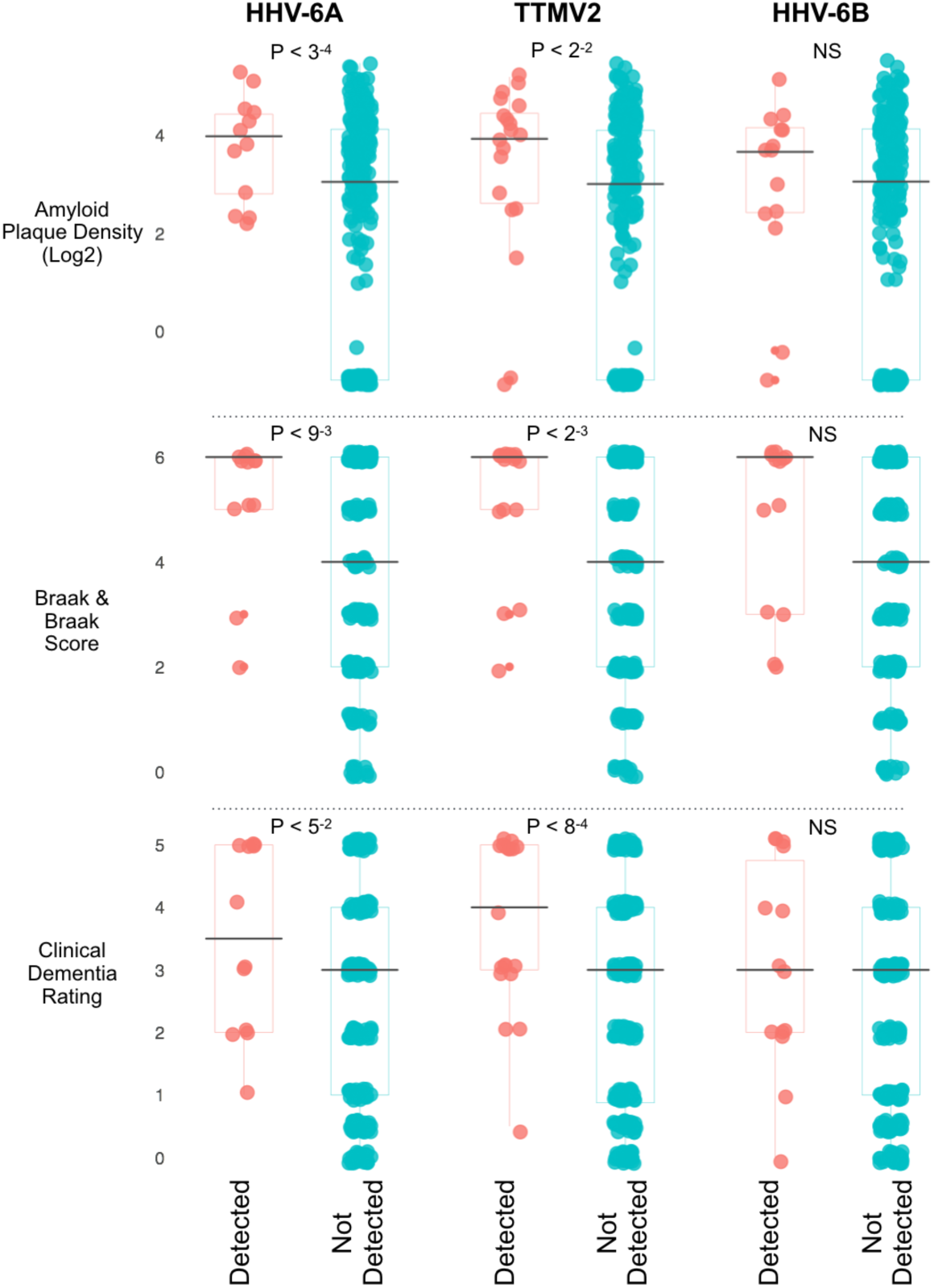
Viral presence association with clinical and pathological traits of Alzheimer’s disease. Boxplots of sample attributes, stratified by presence, or absence of each virus. P-values from two-sample, two-sided T-test. NS: P-value > 0.05.

## Discussion

The generation of large quantities of high quality biological sequence data from the brain tissue of individuals affected by AD offers unprecedented opportunities for building new perspectives into disease biology. The detection and quantification of non-human sequences in NGS data has the potential to contribute to a deeper understanding of the molecular context of AD, but such unanticipated uses of collected data can also present novel methodological challenges.

Although we previously detected a persistently increased abundance of HHV-6A, and HHV-7 across multiple brain regions and across several independent AD cohorts^8^, the quantities of detected virus were generally low compared to the abundance of endogenously expressed genes. Many samples lacked any viral reads. We therefore considered whether the associations that we were reporting^7^ might be associated with spurious signals that did not indicate authentic between-group differences. Using a simulation approach, we observed that low viral abundance and sparsity reduces our power to detect true differences, and correspondingly, is biased against the detection of potentially significant differences. This supports the veracity of our originally reported observation of increased HHV-6A and HHV-7, although also suggests that the viral signal within these datasets is at or near the limits at which a difference might be detected.

We also report new findings of increased viral abundance in AD, based on WGS data generated from subjects within the MSBB. The generally higher quantity of viral reads available in these WGS samples enabled us to use a complementary *de novo* assembly approach to viral sequence quantification that was conceptually distinct to our previous 31mer based method. This offers the opportunity to examine longer nucleotide sequences and to apply strict thresholding to reduce the possibility that technical artefacts or sequence misclassification could inflate viral signals. Consistent with our previous findings, HHV-6A was the virus most consistently associated with AD. As distinct from our previous report, this was driven by a large difference in the fraction of samples in which HHV-6A was detected at all. By examining the categorical presence or absence of HHV-6A, we are able to examine an association between AD and HHV-6A in a manner that does not rely on assumptions necessary for use of conventional differential expression paradigms, and thus provides an encouraging consistency with our previous reports.

We also observed several differences between these additional WGS data and our previous findings. Most notably, in the new WGS dataset, we did not observe an AD-associated increased in HHV-7. We did detect HHV-7 contig clusters within 17 individuals, and although this occurred at a higher rate within AD samples (6.3% AD Definite, 3.3% Normal), this was not statistically significant (P-value: 0.39). Our previous study did include a differential abundance analysis of whole exome sequences (WES) from the MSBB^8^, and, likewise, within these DNA sequences, we also noted an increased abundance of HHV-6A but not HHV-7.

In addition to increased HHV-6A abundance, we found an increased abundance (though not an increased prevalence) of HHV-6B and TTMV2. TTMV2 is a highly prevalent, pan-tropic small anellovirus with unknown relevance to human disease^10^ not detected in our previous study^7^. We did previously report an increased abundance of HHV-6B in MSBB RNA-sequences from the STG and decreased abundance in the APFC.

We observed that, in general, the apparent prevalence of virome signals was lower in the WGS data than in the previously reported RNA sequences. For instance, within the RNA-sequences from the STG, we detected multiple reads mapping to HHV-6A in 32% of samples, whereas in the WGS we detected HHV-6A contig clusters in only 4.3% of samples. This difference is likely partly driven by our choice of methodology for viral quantification. *De novo* construction of viral contigs requires the presence of a relatively large number of candidate reads to assemble, resulting in samples that contain insufficient quantities of viral reads being classified as non-viral. Of those samples yielding signals for HHV-6A contig clusters, the mean number of reads mapping to HHV-6A was 1897 reads, substantially higher than observed in RNAseq from the same region. How these estimates reflect the population prevalence of HHV-6A is not clear, as consistent estimates for these viruses (particularly in the brain) are yet to be clearly established. In a study of HHV-6A and HHV-6B DNA positivity in post-mortem brain samples from five brain regions, Chen *et al^11^* detected HHV-6A in 4% of samples, with signal from at least one brain region for 27.5% of subjects. HHV-6B was detected in 24.3% of samples, with detection in at least one region for 75% of subjects. In a recent study of cerebellar HHV-6A and HHV-6B infection in mood disorders, Prusty *et al^12^* reported the detection of HHV-6A proteins in 22% of control subjects, and HHV-6B in 32%, whereas HHV-6A DNA and protein were detected in only 4%, and HHV-6B DNA and protein was detected in 0% of controls. Lin *et al*^13^ reported that HHV-6A and/or HHV-6B was present in 70% of brain samples from AD subjects, and only 40% of brain samples from age matched controls. Overall, this suggests considerable regional heterogeneity in the prevalence of these *Roseoloviruses* within the brain, and also suggests that our estimated single-region HHV-6A prevalence of 4.3% is a reasonable lower bound of the true prevalence. Although still far higher than reported rates of inherited chromosomally integrated HHV-6 (iciHHV-6) which is estimated at approximately 1% of US and UK control populations^14^, it is plausible that part of our detected HHV-6A signal may be accounted for by iciHHV-6. Further computational and molecular profiling of these samples aimed at resolving this issue are ongoing.

When taken together with our original report^8^, the current study indicates the considerable complexity of the brain virome in AD. In addition to the regional differences in detectable viral signals, we also observed differences in the detectable apparent viral profiles that varied according to methodology employed; i.e., differences across RNA-sequencing, whole exome sequencing, and whole genome sequencing profiling. The extent to which these differences reflect biologically meaningful nuance compared with varying sequence capture and enrichment biases associated with different modalities is also not yet well characterized. Despite the challenges of interpreting platform-associated differences, certain viral signals persist across these sequencing modalities, especially that of HHV-6A. This provides encouraging evidence of the relevance of viral activity to understanding the molecular basis of AD. These observations, when taken together with the findings of many others using a multitude of methodologies^3, 4, 6, 7, 13, 15-21^, support the value of a comprehensive effort to directly profile and characterize the brain microbiome in AD across brain regions, histopathologies, and clinical status.

## Acknowledgments

The results published here are based on data obtained from the AMP-AD Knowledge Portal (doi:10.7303/syn2580853). MSBB WGS data used for the evaluation of viral sequences in AD were generated from postmortem brain tissue collected through the Mount Sinai VA Medical Center Brain Bank and were provided by Dr. Eric Schadt and Dr. Mary Sano from Mount Sinai School of Medicine.

S.G. and M.E.E. acknowledge the support of U01 AG046170 from the NIA. B.R., S.G., M.E.E., and J.T.D. acknowledge the support of 1R56AG058469, 1R01AG058469, and R21AG63968 from the NIA. B.R and J.T.D. acknowledge the support of U01AG061835 and R21AG063068. Philanthropic financial support was provided by Katherine Gehl. The computational resources and staff expertise provided by the Department of Scientific Computing at the Icahn School of Medicine at Mount Sinai also contributed to the performance of this research. The authors also acknowledge Research Computing at Arizona State University for providing computational resources that have contributed to this research.

## Authors’ contributions

B.R., S.G., M.E.E., and J.T.D. designed the study. B.R. performed the computational analysis. B.R., S.G., J.V.H.M., M.E.E., and J.T.D. wrote the paper. All authors read and approved the final manuscript.

## Declarations of Interests

The authors declare that they have no competing financial interests in relation to the work described.

## Supplementary Files

**Table S1:** Summary of viral DNA differential abundance and prevalence analysis

## Materials and Methods

### *De novo* assembly of viral contig clusters

WGS fastq files were downloaded from the Accelerating Medicines Partnership - Alzheimer’s Disease (AMP-AD) Knowledge Portal (synapse ID: syn10901600), performing a preliminary alignment of WGS fastq files to a viral reference database using bowtie2^22^, identifying candidate viral sequences. Mapped reads were filtered through BMtagger^23^ to remove likely human reads. Any putatively non-human reads with low quality scores were trimmed, and reads with a trimmed read length < 60 bases were discarded. Reads were again filtered using BMtagger^23^ to remove likely bacterial reads. Filtered, trimmed, non-human reads were then input to Trinity sequence assembler^24, 25^, including contig formation by the *Inchworm* module, and contig clustering by the *Chrysalis* module. The abundance of each individual contig cluster was estimated by remapping the reads input to Trinity, to the set of contig clusters identified within each sample using bowtie2^22^, using a very sensitive, local alignment, outputting single best alignment to generate a contig cluster / sample count matrix. To identify contig clusters with a likely viral origin, we performed a BLAST^26^ analysis, using BLASTn, to search for any contig clusters with homology to viruses within the original viral database. BLASTn hits with an e-value < 1e-3 matches were classified as viral, and retained for subsequent analysis. Contig clusters with associations to multiple viruses were assigned to a single species based on the highest high-scoring segment pair (HSP). Contig clusters with length > 300 nucleotides, and a mean coverage > 5x were retained for subsequent analysis. Virus level abundance for each sample was estimated as the sum of mapped reads for each contig cluster associated with a particular virus.

### Differential viral abundance analysis

We performed differential viral abundance analysis as previously described^8^. Briefly, definitions of AD were based on the multiple levels of CERAD neuropathology classification^27^. Within each comparison, we retained viruses with multiple mapped reads in at least 10 samples. Viral counts were adjusted for the total number of reads in each fastq file, and normalized using the Voom function in the Limma package^9, 28^. Linear models were fit for each virus, including covariates for: AD status, age of death, sex, ethnicity, and post-mortem interval (PMI). Differential abundance between AD and Normal groups were estimated using the eBayes function^29^, setting robust = TRUE to minimize the effect of outliers in variance.

### Differential viral presence analysis

Within each comparison, we retained viruses with multiple mapped reads in at least 10 samples. For each virus, we identified samples with, and without detected virus, and used a two-sided Fisher’s exact test to detect unexpectedly high, or low overlap between virus-associated samples, and AD samples.

### Viral association with clinicopathological traits of Alzheimer’s analysis

For each AD trait under consideration (Clinical Dementia Rating; Braak score; mean amyloid plaque density), we iterated each feature across each virus detected within at least 3 samples. We compared the distributions of each AD trait between virus-associated and non-virus-associated samples using a two-sided, two-sample T-test. Given that mean amyloid plaque density demonstrates a right skewed distribution, we first incremented scores by 0.1, followed by log2 transformation to normalize the distribution prior to testing.

